# A Porcine Model of Intervertebral Disc Injury Recapitulates Human Discogenic Pain via Notochordal Cell Loss and Pain-inducing Nucleus Pulposus Cell Emergence

**DOI:** 10.1101/2025.11.24.690318

**Authors:** Giselle Kaneda, Jacob T. Wechsler, Melissa Chavez, Julia Sheyn, Karandeep Cheema, Chushu Shen, Dante Rigo De Righi, Pablo Avalos, Yibin Xie, Wafa Tawackoli, Candace Floyd, Debiao Li, Dmitriy Sheyn

**Affiliations:** Orthopaedic Stem Cell Research Laboratory, Cedars-Sinai Medical Center, Los Angeles, CA; Board of Governors Regenerative Medicine Institute, Cedars-Sinai Medical Center, Los Angeles, CA; Department of Biomedical Sciences, Cedars-Sinai Medical Center, Cedars-Sinai Medical Center, Los Angeles, CA; Biomedical Imaging Research Institute, Cedars-Sinai Medical Center, Los Angeles, CA; Department of Bioengineering, University of California, Los Angeles, CA; Department of Orthopaedics, Cedars-Sinai Medical Center, Los Angeles, CA; Department of Surgery, Cedars-Sinai Medical Center, Los Angeles, CA; Department of Emergency Medicine, Emory University, Atlanta, GA

**Keywords:** Discogenic pain, Low back pain, Porcine model, Intervertebral disc degeneration, Biobehavioral tests, Notochordal cells, Nucleus pulposus

## Abstract

Lower back pain (LBP) is one of the most common causes of disability, with up to 40% of LBP cases attributed to intervertebral disc (IVD) degeneration. While small animal models are widely used to study IVD and LBP, the small size of their IVDs limits translational and biological relevance. Large animal models more accurately emulate human disease; however, methods of measuring LBP are not well established. Pigs were also considered unfit for LBP research, due to notochordal cells (NCs) persistence through life, unlike humans. We developed a comprehensive porcine model with quantitative measure of discogenic pain via biobehavioral testing (BBT), MRI, and multi-omics tissue analysis of IVD and DRGs. MRI demonstrated the progression of IVD degeneration beginning at 4 weeks post-injury. BBTs showed the development of significant pain responses by week 4 post-injury, supported by transcriptomics of injury matched DRGs. Single cell transcriptomics, trajectory and cell-cell communication analyses suggest that, with injury, NCs are differentiating to nucleus pulposus cells (NPCs). Furthermore, NPCs showed upregulation of cellular stress, neural outgrowth, and inflammation pathway, consistent with pain-inducing distress signals found in human samples. This study establishes novel MRI and BBT-based methods for quantifying LBP in pigs and supports its translational relevance to human discogenic LBP. The identification of LBP-associated clusters mirrors our previous finding in humans. Moreover, the shift of NC to NPC phenotype further supports that the porcine model is relevant to human pathology, as the injury induced accelerated aging and loss of NCs with IVD degeneration and discogenic pain.

## Introduction

Lower back pain (LBP) is the greatest cause of disability worldwide with an estimated 619 million people suffering globally [20]. This number is expected to rise to 843 million by 2050 due to population aging [20]. Up to 40% of LBP cases can be attributed to intervertebral disc (IVD) degeneration [23; 78]. However, the relationship between disc degeneration and pain is complex, with many cases of disc degeneration being asymptomatic. The disconnect between structural pathology and clinical symptoms has resulted in therapeutic development for LBP treatments falling behind other conditions. Current treatments focus on symptom management and include pharmacological therapies, such as non-steroidal anti-inflammatory drugs or opioids, and invasive surgical procedures [24]. Thus, there is a clinical need for LBP treatments that alleviate symptoms and reverse disc degeneration.

Discogenic LBP is a multifactorial disease with multiple factors implicated in its development including vascularization, inflammation, and innervation [62]. The mechanisms that underpin LBP development have yet to be fully elucidated. The IVD is composed of 3 distinct tissues: the cartilaginous endplate, which adheres the IVD to the adjacent vertebral bodies; the annulus fibrosis (AF), which is the outer portion of the IVD made of concentric rings of fibrocartilage; and the nucleus pulposus (NP), the jelly-like center of the IVD that develops from the embryonic notochord [62]. In the first decade of life, humans lose a majority of their notochordal cells [44], and NP cells (NPCs) have little regenerative capacity [12]. During the progression of IVD degeneration, insult to the IVD can induce cell stress and changes to the mechanical and biochemical homeostasis of the IVD. Stressed cells can induce nociceptive and pro-inflammatory signaling, thus triggering pain and inflammatory cascades within the IVD, further altering IVD composition [43]. With sustained degeneration, NPCs begin to die off, further exacerbating disruptions to the IVD micro-environment [101].

In vivo LBP studies are primarily performed in small animal models due to the low cost [34] and well-established biobehavioral testing methods that can quantify various forms of pain. However, small animal models lack the translational relevance to advance preclinical treatments into clinical trials. Specifically, the smaller IVD size in rodents results in a significant differences in nutrient diffusion patterns, biomechanical loading, and cellular microenvironment compared to human IVD, limiting the relevance of findings to human pathophysiology [22]. In comparison, large animal models like pigs offer superior translational potential due to their closer anatomical similarities to humans, including disc size and nociceptive pathway organization [58; 84].

In this study, we have comprehensively analyzed a [8; 61; 79; 105] large animal model of IVD degeneration and LBP in pigs. We have utilized our previously established qCEST MR imaging sequences [8; 17; 68; 79; 105] with biobehavioral testing (BBT) to quantify pain. We characterized the cellular and molecular landscape of injured IVD with injury and confirmed the model’s relevance to human discogenic LBP despite anatomical differences such as the presence of NCs in healthy porcine discs. Finally, we observed the development of pain-inducing cells, reflecting what has been observed in human samples [43].

## Methods

### IVD degeneration animal model

Appropriate Institutional Animal Care and Use Committee (IACUC) approval (protocol No. 008066) was obtained prior to conducting any experiments. Due to safety concerns, namely aggression and handling difficulties associated with boars, only female pigs were used in this study to ensure staff safety and experimental feasibility. Survival surgery was performed on 9 female 4–5-month-old Yucatan minipigs (Premier Biosource, Ramona, CA). Pigs were housed on a 12/12-hour light-dark cycle and were fed once a day in the morning with 32oz of feed.

Sedation was achieved using an intramuscular combination of Ketamine (37mg/kg) and Midazolam (0.1-1 mg/kg) or Telazol (4-6mg/kg). Anesthesia was induced via intravenous administration of Propofol (1-2mg/kg) and maintained with 2-3% inhaled isoflurane via an endotracheal tube with mechanical ventilation. To monitor vital signs, equipment including but not limited to pulse oximeter, capnograph, rectal thermometer, non-invasive blood pressure, and EKG were attached to the animal for the duration of anesthesia. The animals lower thoracic and lumbar regions were prepped, using a disinfectant scrub preparation (e.g., betadine or chlorhexidine) and scrubbed for a minimum of three minutes. Any foam was removed with a clean wet towel, and the skin further disinfected with alternating betadine and 70% isopropyl alcohol wipes three times each. Prior to beginning surgery, pigs received Carprofen (0.5mg/kg) and Buprenophine (0.01-0.02mg/kg) intramuscularly for analgesia. Under fluoroscopic guidance the L3-L4, L4-L5, and L5-L6 were identified and percutaneously punctured with a 17G needle (Fig. 1A-B). Once entry into the disc was confirmed in two orthogonal planes of view, the needle was rotated 360 degrees three times and slowly removed. Post-operatively, the pigs received Carprofen (0.5 mg/kg) every 24 hours for 48 hours and Buprenorphine (0.01-0.02 mg/kg) every 12 hours for 24 hours, both administered intramuscularly.

**Figure 1:**
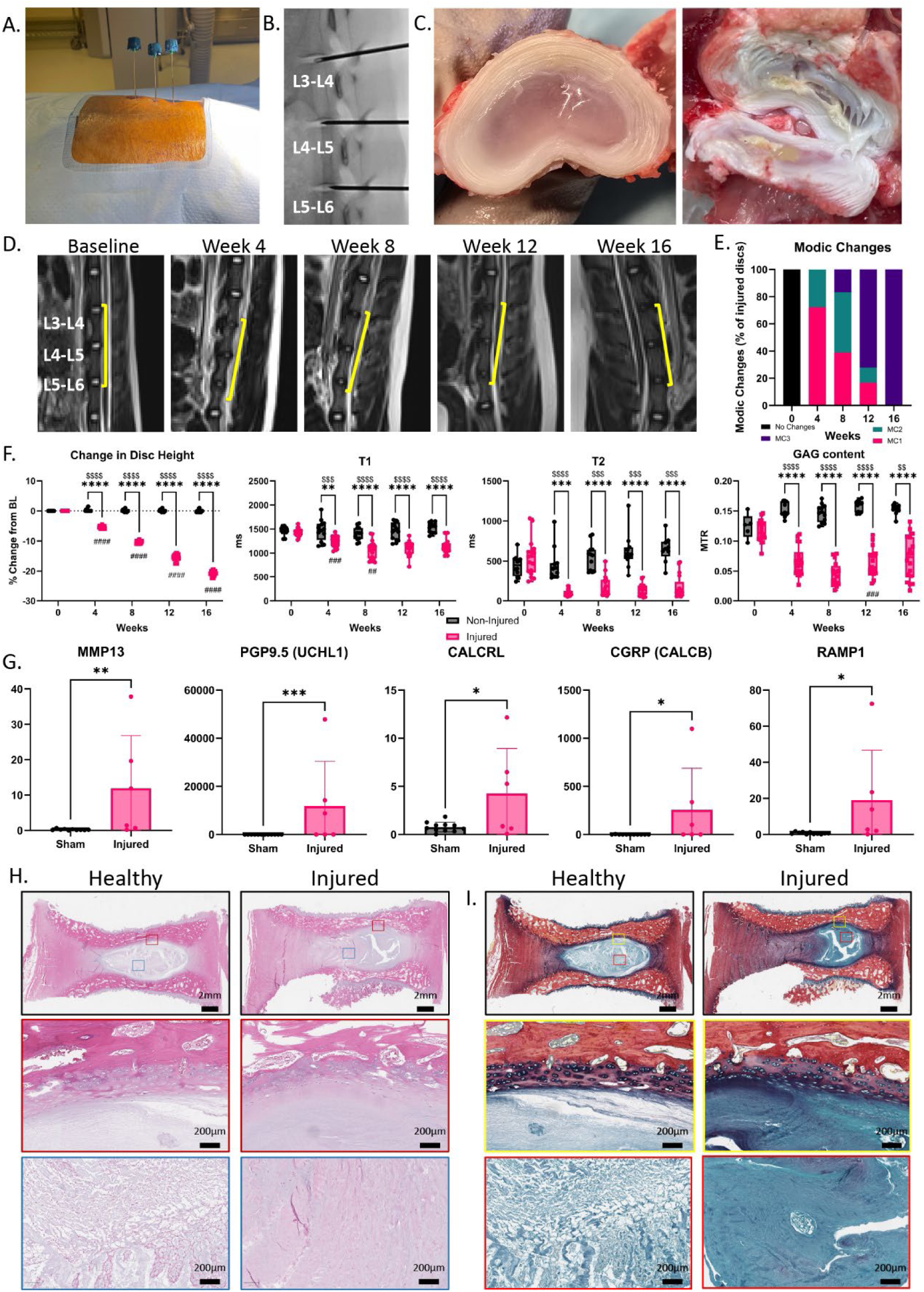
Disc puncture induces progressive degeneration monitored with MRI, molecular, and histological hallmarks of disease. (A) Injury induction, (B) Fluoroscopic view of disc puncture, (C) macroscopic view of non-injured (left) and injured (right) IVD at week 12, (D) Representative images of T2 MRI, (E) percent of adjacent vertebrae with each Modic changes, (F) Gene expression analysis of the IVD for select degeneration, innervation, and CGRP receptor pathway markers; histological staining of non-injured and injured IVD with (G) H&E and (H) Alcian blue/pircrosirius red. *p<0.05,** p<0.01,*** p<0.001,****p<0.0001 between groups. ^$$$^p<0.001, ^$$$$^p<0.0001, injured compared to baseline. ^##^ p<0.01, ^###^ p<0.001, ^####^ p<0.0001 injured compared to previous timepoint.

For euthanasia, pigs were sedated with intramuscular Telazol (4-6mg/kg). Anesthesia was induced via intravenous administration of propofol (1-2mg/kg) followed by 1mL per 4.5kg of Euthasol (Virbac, Westlake, TX) once anesthesia was achieved.

Sham DRG and NP samples for gene expression were generously donated by Dr. Eugenio Cingolani from pigs undergoing terminal cardiac procedures.

### Biobehavioral Testing (BBT)

Biobehavioral testing (BBT, n = 6) was utilized to quantify the development of pain following IVD injury. Tests were adapted from established porcine spinal cord injury models and selected to assess different components of the pain phenotype.

Prior to injury, pigs underwent four weeks of socialization and tester training to minimize adverse behaviors and stress-related responses. One week prior to injury and every other week post injury, pigs underwent a full battery of BBTs.

Testing was performed during morning feeding to minimize behavioral noise and interest in the tester and test administration. Prior to testing, co-housed pigs were separated into individual kennels and randomly taken to the testing room and tested one at a time. Pigs remained in the testing room for approximately 25 minutes, with 10 minutes of acclimation to the testing room, followed by 15 minutes of testing. Following completion of the test, pigs were returned to single housing with any remaining food. After all pigs were evaluated, they were returned to their original housing conditions.

### Glasgow Pain Score

The Glasgow pain scale was adapted from a goat model to measure acute pain.[81] Each pig was assessed daily before feeding and BBT on five criteria as noted on Table 1. All the scores collected in a single testing week were averaged for a final mean week score.

### Wind-up ratio (WUR)

The wind-up ratio test assessed temporal pain summation. Each pig was led to the testing room one at a time. Following acclimation to the room, pigs underwent testing in a control area innervated by non-injured DRGs located between the pig’s shoulder blades, and an injury matched DRGs innervated “testing area” adjacent to the spine just above the iliac crest. Pigs were enclosed in a testing area with their full food bowl placed in the middle of the arena.

A pin prick stimulus was applied to the target area for 2 second, followed by 2 seconds off, repeated 10 times. The number of stimuli with a positive response is recorded. After completing 10 pin pricks in the target area, the control area is completed. Three repetitions were performed per area, alternating between the two.

### Pig Evoked Pain score

Pain-related responses to sensory stimuli were quantified using a novel instrument we developed, the Pig Evoked Pain score (PEPS), which integrates both reflexive (spinal) and cognitive/affective (supraspinal) behavioral responses. Each stimulus was assigned a score from 0 (no response) to 10 (extreme pain).

Supraspinal responses to stimuli included vocalizations, interruption of eating behavior, and escape behavior. Vocalizations were included as an affective indicator based on evidence that pigs, as socially complex animals, use vocal communication to convey meaningful information during distressing experiences [2; 67; 94]. Interruption of eating behavior was adapted from human clinical pain research and was used as an ethological correlate of pain-related activity disruption [32]. To avoid potential confounding effects of hunger on pain perception, pigs were provided a daily ration during testing [39]. Escape behavior was included based on its established association with high-intensity nociceptive stimuli in pigs [50; 87–89] and was scored toward the higher end of the PEPS scale. Reflexive responses were evaluated based on limb withdrawal and whole-body withdrawal from the stimuli, as performed in rodent pain models [25].

PEPS was utilized to quantify pain intensity during WUR testing. Video and audio recordings collected during each session were analyzed post-hoc to assign a PEPS using behavioral ethogram by investigators naïve to the treatment condition, as previously described. Each stimulus was independently assessed by two trained observers who reviewed all available camera angles. Individual PEPS were averaged to produce a final score for each stimulus.

### MR imaging

Prior to MRI, pigs were sedated with intramuscular Telazol (4-6mg/kg). Anesthesia was induced via intravenous administration of propofol (1-2mg/kg) and maintained with 2-3% inhaled isoflurane via an endotracheal tube.

MRI was performed at baseline (pre-injury) and every four weeks post injury, up to 16 weeks post injury on the lumbar spine (L1-L6 levels, n=3 injured IVD and n=2 non-injured IVD). Imaging included traditional (T1 and T2) and novel qCEST sequences to assess disc degeneration and pain development as previously described.[8; 68; 105] In addition, mean disc height was quantified from axial T2-weighted images by calculating the average disc height at each timepoint and level. Adjacent vertebrae (n=18) were independently assessed for Modic changes by two experienced radiologists, using standardized grading criteria adapted from Fields et al.[27] Each disc was scored on a scale of 0–3 with 0 indicating no signal change in vertebral bone marrow and 3 indicating hyperintensity suggestive of severe endplate degeneration.

### Blood collection and processing

While anesthetized for surgery and imaging, two 5mL blood samples were collected: one in a serum separator tube (SST) for serum isolation and one in sodium heparin-coated tube for plasma collection. Following collection, the blood was centrifuged at 2000G for 10 minutes and the resulting serum and plasma aliquoted and stored at -80°C for later analysis.

### Cell Preparation for Single Cell RNA Sequencing

The NP and surrounding transition zone (n = 3 injured and 3 non-injured discs) were isolated and manually morselized to ∼1mm^3 pieces. The tissue was then enzymatically digested for 1 hour at 37°C in 2 mg/mL of Pronase Protease (Millipore, Temecula, CA) in growth media containing 1% antibiotic-antimycotic solution (Gibco, Carlsbad, CA) and 10% fetal bovine serum (FBS, Gemini Bioproducts, West Sacramento, CA) in Dulbecco’s modified eagle media-F12 media (DMEM/F12, GIBCO, Carlsbad, CA), followed by an ∼18 hours digestion at 37°C in 0.25 mg/mL of Collagenase Type 1S (Sigma Aldrich) in growth media. The resulting sample was pushed through a 70μm cell strainer, and cells were isolated via centrifugation at 300G for ten minutes. The pellet was resuspended in phosphate-buffered saline at a concentration of 1,500 cells/uL.

### scRNA-seq Library Preparation and Sequencing

Single cell suspensions were stained with ViaStain AOPI Staining Solution (Nexcelom Bioscience, Lawrence, MA) and imaged on an EVOS M7000 Imaging System (Thermo Fisher Scientific, Waltham, CA) to determine cell suspension quality and cell viability. Cell suspensions with a minimum viability of 70% were partitioned (10,000 target capture) into Gel Beads-in-emulsion (GEMs) on the Chromium X (10x Genomics, Pleasanton, CA) per the Chromium NextGEM Single Cell 3’ v3.1 Gene Expression kit user guide. Following reverse transcription and cDNA amplification, single cell libraries were prepared on the Chromium Connect (10x Genomics) using the Automated Gene Expression Library Construction kit (10x Genomics). Barcoded libraries were quantified by quantitative PCR using the Collibri Library Quantification Kit (Thermo Fisher Scientific) and library size was measured via the 4200 TapeStation (Agilent Technologies, Santa Clara, CA). Libraries were sequenced on a NovaSeq 6000 or X Plus (Illumina, San Diego, CA) with sequencing configuration 28x10x10x90bp, and a sequencing depth of 50,000 reads/cell.

### scRNA-seq Bioinformatic Analysis

Raw sequencing data was demultiplexed and converted to FASTQ format using bcl2fastq v2.20. Cell Ranger v7.1.0 (10X Genomics) was used for barcode identification, read alignment, and UMI quantification with default parameters and aligning to the pig reference genome Sscrofa11 with intron mode used.

Following initial processing, the samples were transferred to R (v4.3.3) for downstream analysis. Briefly samples were converted from a 10X filtered count matrix to a Seurat object (Seurat package, v5.1.0) using the *CreateSeuratObject* function.[38] At this time, genes expressed in less than three cells, and all cells expressing less than 200 genes were removed from each Seurat objects. Next, samples were normalized using the *NormalizeData* function and the top 8000 most differentially expressed genes identified using *FindVariableFeatures* with the *“vst”* method. To prepare for integration, shared features were identified using *SelectIntegrationFeatures*. Afterward, *FindIntegrationAnchors* and *IntegrateData* were used to integrate and anchor the 6 samples’ data. *RunPCA* and *RunUMAP* functions were used for principal component analysis with PC=15. *DimPlot* was used with the *“pca”* reduction to visualize sample-specific distributions and integration quality. *FindNeighbors* and *FindClusters* functions were then used for unsupervised clustering at a resolution of 0.9. The RNA assay was set as the default for downstream analyses, and data layers were unified using *JoinLayers*. Cluster identities were identified using canonical markers gene: TBXT, KRT8, KRT18, and CD24 for NCs; MGP, DCN, COMP, and SERPINE2 for NPCs; and PECAM1, ENG, and CDH5 for EC (Supplemental fig 3). Clusters co-expressing both NC and NP markers were classified as transitional (NC-NP, Supplemental fig 3). Proliferative/mitotic clusters were identified using the marker TOP2A, CDK1, and BIRC5 (Supplemental fig 3).

### Trajectory analysis (Monocle 3)

Trajectory analysis was performed using the Monocle3 (v1.3.1) package.[15; 36; 52; 70; 85; 86] The combined Seurat object was converted to a cell_dataset_set using the *as.cell_data_set* function and run through the Monocle 3 pipeline. Briefly, the object was normalization, highly variable genes selected, and dimensionally reduce via the *preprocessing_cds* function. The top 100 prinicipal component were stored for downstream analysis. Next, the *align_cds* function was used to batch corrected and … Disc identity as injured or healthy was used for the alignment group and UMAP was used for the reduction method. [60] Next, dimensionality was further reduced and the cells clustered using the *reduce_dimension* and *cluster_cells* functions. The trajectory root was programmatically determined using the *get_earliest_principal_node* function.

### Cell-cell communication analysis (CellChat)

Cell-cell communication analysis was performed using the CellChat (v2.1.2) package.[46] Briefly, the integrated Seurat object was converted to a CellChat object using the *createCellChat* function. Due the lack of a pig specific database, the human database CellChatDB.human was used. Data was subset to include only relevant data via the *subsetData* function then overexpressed genes and ligand-receptors identified using *identifyOverExpressedGenes and identifyOverExpressedInteractions.* To predict expressed pathways, the function *computeCommunProbPathway* and *aggregateNet* were used.

### ELISA

Serum protein content was quantified using a Pierce BCA kit according to manufacturer protocol (Thermo Fisher, Walthan, MA). Serum IHH content was measured using IHH ELISA (MyBioSource, San Diego, CA) following manufacturer protocol. Absorbance was measured using a SpectraMax M3 (Molecular Devices, San Jose, CA).

### Gene Expression

Total RNA for the nucleus pulposus and dorsal root ganglia were collected using a RNeasy Lipid Tissue Mini Kit (Qiagen, Hilden, Germany) following the manufacturer protocol. The isolated RNA was transcribed into cDNA using a high-capacity reverse transcription kit (Applied Biosystems, Waltham, MA). Pig Taqman gene expression assays (Thermo Fisher Scientific) as detailed in Supplemental materials were used for analysis (Supplemental Table 2). All assays were normalized to 18S housekeeping gene.

### DRG Total RNA-seq library preparation and sequencing

RNA integrity was analyzed on the 2100 Bioanalyzer using the Agilent RNA 6000 Nano Kit (Agilent Technologies, Santa Clara, CA) and RNA quantified using the Qubit RNA HS Assay Kit (ThermoFisher Scientific, Waltham, MA). Four hundred ng RNA was ribodepleted using the Lexogen RiboCop Depletion Kit Human/Mouse/Rat v2 kit (Lexogen Inc., Greenland, NH). Stranded RNA-Seq library construction was performed using the IDT xGen Broad-Range RNA Library Prep Kit (Integrated DNA Technologies, Coralville, IA) with 11 cycles of PCR amplification. Library concentration was measured via the Qubit 1X dsDNA HS Assay kit, and library size on the Agilent 4200 TapeStation (Agilent Technologies) with the Agilent HS D1000 ScreenTape. Multiplexed libraries were pooled and sequenced on a NovaSeq 6000 (Illumina, San Diego, CA) using 1x75bp sequencing at 50M reads/sample.

### IVD Total RNA-seq library preparation and sequencing

RNA integrity was analyzed on the 2100 Bioanalyzer using the Agilent RNA 6000 Pico Kit (Agilent Technologies, Santa Clara, CA) and RNA quantified using the Qubit RNA HS Assay Kit (ThermoFisher Scientific, Waltham, MA). Library construction was performed using the SMARTer Stranded Total RNA-Seq Kit v3 – Pico Input Mammalian kit (Takara Bio USA, Inc., Mountain View, CA). Three ng of total RNA was used for library construction with 13 cycles of PCR amplification. Final libraries were quantified via the Qubit 1X dsDNA HS Assay kit and analyzed for size on the 4200 TapeStation (Agilent Technologies) with the Agilent HS D1000 ScreenTape. Multiplexed libraries were pooled and sequenced on a NovaSeq 6000 (Illumina, San Diego, CA) using 1x75bp sequencing at 50M reads/sample.

### Total RNA-seq Data analysis and processing

Raw reads obtained from RNA-Seq were aligned to the transcriptome using STAR (version 2.6.1) (Dobin A et al., 2013) / RSEM (version 1.2.28) (Li B and Dewey CN, 2011) with default parameters, using a custom pig Sscrofa11 transcriptome reference downloaded from https://useast.ensembl.org/, containing all protein-coding and long non-coding RNA genes based on Sscrofa11.1.104 annotation.

### Proteomic Sample Preparation

S-Trap high recovery lysis buffer (5% SDS, 8M Urea, 100mM glycine) was added to approximately 10mg of macerate NP tissue or plasma. The mixture was incubated for 30 minutes on ice and then sonicated at 4°C using a Bioruptor Pico Sonicator (Diagenode, Denville, NJ) for 10 minutes at 30 seconds on and 30 seconds off. Lysate was then centrifuged at 12,000G for 10 minutes at 4°C, and the supernatant was saved for downstream processing. 100ug of protein lysate was used for downstream processing. Briefly, 1,4-dithiothreitol (Sigma Aldrich) was added to the sample for a final concentration of 20mM of DDT and heated for 15 minutes at 37°C. Next, iodoacetamide (Sigma, Aldrich) was added to the solution for a final concentration of 40mM and incubated in the dark for 30 minutes. Then, phosphoric acid was added for a final concentration of 1.2% phosphoric acid. At this point, pH was checked using a pH strip to ensure sample pH was between 1-2 to ensure proper binding of the protein to the spin column. The sample was added to a S-Trap (Protifi, Fairport, NY) and washed with S-Trap binding buffer (100mM TEAB in 90% MeOH). Samples were digested for one hour at 47°C with 1ug of trypsin per 25ug of protein lysate. Following digestion, samples was eluted using 50mM TEAB, 0.1% Formic Acid in LC-MS grade water, followed by a 1:1 solution of 0.1% Formic Acid in LC-MS grade water and acetonitrile. Samples were presumed to have lost approximately 50% of their original protein quantity during the S-Trap digestion. Eluted samples were dried down via SpeedVac vacuum concentrator (Thermo Fisher, Waltham, MA), then resuspended in 0.1% FA in LC-MS grade water for a final concentration of 0.2ug/uL.

### LC-MS/MS Data Acquisition and Analysis

DIA analysis was performed on an Orbitrap Ascend Tribrid (Thermo Scientific) mass spectrometer interfaced with an EASY-Spray™ ionization source (Thermo Scientific, ES081) coupled to Vanquish Neo ultra-high-pressure chromatography system with 0.1% formic acid in water as mobile phase A and 0.1% formic acid in acetonitrile as mobile phase B.

Proteomic (PTM) peptides were separated at constant flow rate of 1.20 µL/minute with a linearly increasing gradient of 8-34% B for 0-90 minutes, then flushed with 98% B from 90-112 minutes. The column used was Thermo Scientific™ µPac™ HPLC column with a 200cm bed length (P/N: COL-NANO200G1B). MS1 resolution was set to 120,000 with AGC target set to standard. RF Lens was set to 60% with a maximum injection time of 251 ms. Fragmented ions were detected across a scan range of 380-985 m/z with 60 non-overlapping data independent acquisition precursor windows of size 10 Da. MS2 resolution was set to 15,000 with a scan range of 145-1450 m/z, a normalized collision energy of 30%, and a normalized AGC target of 400% with a custom maximum injection time set to 25 ms. All data is acquired in profile mode using positive polarity.

For IVD cell and tissue samples acquisition, field-assisted ion mobility spectrometry (FAIMS) was enabled using a single compensation voltage (CV) of –45. Plasma samples were acquired without FAIMS under otherwise identical conditions.

Proteins were identified from raw MS files and quantified using DIANN 1.8.1 against spectral library generated from the NCBI Sus scrofa 11.1 reference genome, using matches between runs and reannotation of gene groups to a supplied porcine FASTA protein sequence database, but with no heuristic protein inference and no normalization applied. Protein quantification estimates were produced using the DIANN maxLFQ algorithm, and only proteins quantified by proteotypic peptides were included in the final dataset for further analysis.

Bioinformatic analysis of the was performed using the MetaboAnalyst web platform v6.0. Prior to analysis, any duplicate proteins or peptides with more than one protein identification were removed. Data was then normalized by median. A minimum absolute fold change (FC) of 2-fold difference and adjusted false discovery rate of p < 0.1 were selected as a significant.

### Histological and Immunofluorescent Staining of IVD

IVD were harvested for histology and immunofluorescence (n = 4 per group). Following sacrifice, the IVD and part of the vertebral body were dissected and fixed in 10% phosphate buffered formalin for 72 hours. The IVD were then decalcified in 6% TCA for 4 weeks, passed through a graded series of ethanol solutions, and embedded in paraffin. Five-micron-thick sections were cut from the paraffin blocks. Hematoxylin and eosin (H&E) and Alcian blue/Picrosirius red (AB/PR) staining were performed to evaluate morphological features of healing, tissue organization, and scar tissue formation. All histological slides were imaged at 20x using a Leica Aperio AT2 slide scanner.

For immunofluorescent (IF) staining, tissues were deparaffinized, and the antigens were retrieved by incubation in Proteinase K (Agilent, Carpinteria, CA) for 12 minutes at room temperature. Nonspecific antigens were blocked by applying serum-free protein block (Agilent). Slides were stained with primary antibodies, as detailed in Supplemental Table 2. The primary antibodies were applied to the slides, after which the slides were incubated at 4°C overnight and washed using PBS; the slides were then incubated with secondary antibodies for one hour at room temperature. Finally, the slides were stained with DAPI for five minutes in the dark. Fluoromount-G (Invitrogen) was applied on the tissue and the tissue coverslipped. Images were captured using an Axio Imager Z1 fluorescent microscope (Carl Zeiss, Oberkochen, Germany) equipped with ApoTome and AxioCam HRc cameras. Negative controls were processed using identical protocols while omitting the primary antibody to exclude nonspecific staining.

### Immunofluorescent Staining of DRGs

DRGs were harvested for immunofluorescence (n = 2 per group). Following sacrifice, the DRGs were identified and snap frozen in liquid nitrogen. The DRG was placed into OCT medium for cryosectioning. DRGs were cut at a 20um thickness and fixed using 4% PFA for 15 minutes. Antigens were retrieved by incubation in Proteinase K (Agilent) for 12 minutes at room temperature. Nonspecific antigens were blocked by applying serum-free protein block (Agilent). Slides were stained with primary antibodies, as detailed in Supplemental Table 3. The primary antibodies were applied to the slides and incubated at room temperature overnight. Following incubation, slides were washed using PBS and incubated with secondary antibodies for one hour at room temperature. Finally, the slides were stained with DAPI for five minutes in the dark. Fluoromount-G (Invitrogen) was applied to the tissue and the tissue coverslipped. Images were captured using a Leica Stellaris 8-STED Super-resolution Confocal Microscope and Axio Imager Z1 fluorescent microscope (Carl Zeiss, Oberkochen, Germany) equipped with ApoTome and AxioCam HRc cameras. Negative controls were processed using identical protocols while omitting the primary antibody to exclude nonspecific staining.

### Statistical Analysis

All statistical analysis for BBT, MRI measurements, and gene expression were performed in GraphPad Prism 10 (GraphPad, La Jolla, CA). Outliers greater than two standard deviations from the mean were excluded.

For BBT, MRI measurements, and gene expression of DRGs, 2-way analysis of variance (ANOVA) or mixed-effects analysis, were performed for each dependent measure separately, using mean values with grouping of experimental groups. For multiple comparisons, appropriate post hoc tests were used.

For gene expression analysis of IVD, normality was assessed using the Shapiro-Wilk test. Normally distributed samples were compared using unpaired t-tests while non-normally distributed samples were compared using Mann-Whitney U tests.

## Results

### Confirmation of IVD degeneration following annular puncture

Gross comparison of injured and non-injured IVD showed reduced NP size and loss gelatinous consistency, at 12 weeks post injury (Fig. 1C). MR imaging of the injured and adjacent non-injured IVD showed significant changes in IVD morphology starting at 4 weeks post injury (Fig. 1D-F). Type I and II Modic changes in the adjacent vertebral were detected at week 4 post injury, with all adjacent vertebra demonstrating Type III changes by 16 weeks post injury (Fig. 1E). Significant reductions in disc height were observed between injured and non-injured IVDs (significance indicated by *), injured compared to baseline (significance indicated by $), and with the previous timepoints (significance indicated by #) at all timepoints post injury (Fig. 1F). Longitudinal and transverse relaxation time, represented by T1 and T2 respectively, and MTR which indirectly quantifies glycosaminoglycan content showed similar trends (Fig. 1F).

Gene expression analysis of the IVDs at 16 weeks post injury demonstrated significant upregulation of select degeneration (MMP3), neuronal innervation (PGP9.5 [5; 66]), and CGRP receptor pathway (CALCRL [41; 69; 99], CGRP [41; 69; 99], RAMP1[41; 69]) genes in injured IVD compared to sham discs (Fig. 1G). Select markers for inflammation (IL1β [71], TNFα [48]) as well as NGF related (NGF [6; 31] , TrkA [40]) and nociceptive peptide/channel markers (TRPV1 [30], TAC1 [33; 107]) demonstrated a similar trend of increased expression with injury (Supplemental Fig. 1).

Histological staining with H&E and AB/PR shows a marked loss of ECM organization with accumulation of dense ECM consistent with fibrotic remodeling and scar tissue replacement (Fig. 1H and 1I).

### Evaluation and validation of chronic pain development

MR imaging using qCEST sequence was previously validated in human patients [56; 68] and correlated to low pH in a pig model [8; 105]. Therefore, we have used this method to validate pain outcome measures in this study. Here the qCEST indicated a significant decrease in pH starting at week 8 and continuing until euthanasia at week 16 in injured IVD compared to non-injured controls (Fig. 2A). Acute pain as assessed by the Glasgow pain scale was found to be significantly higher compared to baseline around week 10 and continuing to increase up until euthanasia (Fig. 2B). Temporal summation of pain, as measured by Wind Up ratio (WUR) was found to be significantly increased in injury matched dermatomes starting at week 4 and continuing up to 14 weeks post injury (Fig. 2C). Furthermore, the highest pig evoked pain response (PEPS) score was found to be significantly higher in the injury matched dermatomes compared to the control dermatome (Fig. 2D).

**Figure 2:**
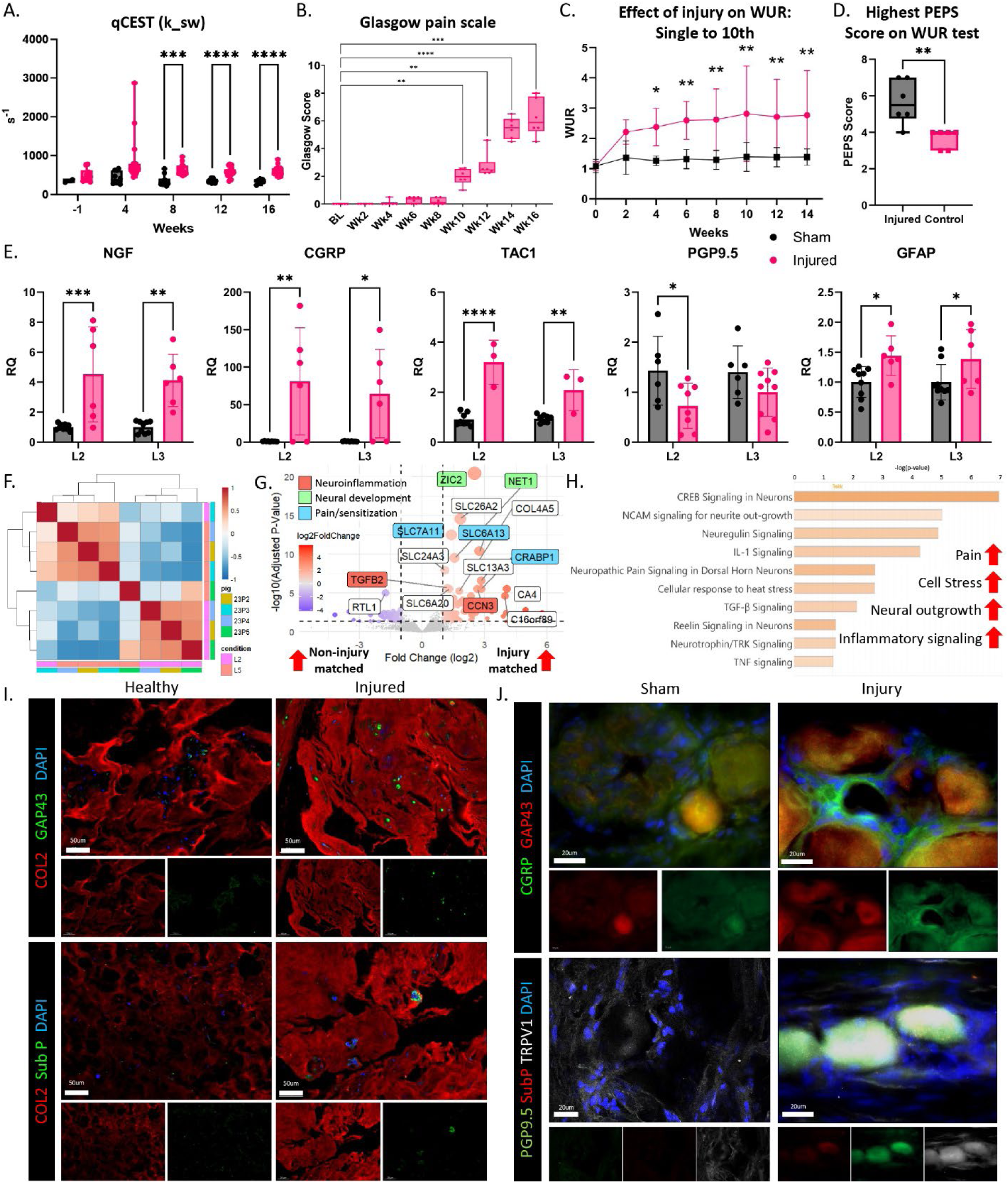
Injury triggers pain behaviors in addition to transcriptional and cellular changes in DRGs. (A) qCEST MR imaging, (B) Glasgow pain scale, (C) Wind Up Ratio, (D) highest PEPS score on Wind Up Ratio, (E) Gene expression analysis of injury match and sham DRGs for select pain and neuroinflammation markers; (F) Heatmap of injured matched and non-injury matched DRGs, (G) volcano plot of top 15 DEGs from healthy vs. injured DRGs, (H) IPA pathway analysis of select significantly enriched pathways, IF staining of (H) IVD and (I) DRGs.

Gene expression analysis of pain (NGF [6; 31] , CGRP [41; 69; 99], TAC1 [33; 107], PGP9.5 [5; 66]) and neuroinflammation (GFAP [37; 42; 96]) in sham and injury match DRGs demonstrated significantly higher pain and neuroinflammation in the injury matched DRGs compared to level matched shams (Fig. 2E). Further comparison of the injured (L2 and L3) and non-injured matched levels (L4 and L5) demonstrated that upregulation occurred only at the level of injury (Supplemental Fig. 2). Bulk transcriptomics of DRGs from injury matched and non-injury matched dermatomes demonstrate distinct transcriptional profiles (Fig. 2F). Multiple genes associated with neuroinflammation (CCN3 [49], TGFβ-2 [16]), neural development (NET1 [92], ZIC2 [63]), and pain/sensitization (CRABP1 [64], SLC6A13 [35; 100], SLC7A11 [7]) were found to be enriched in the injured DRGs (Fig. 2G). IPA pathway analysis predicted upregulation of pathways associated with pain, cell stress, inflammation, apoptotic signaling, and neural outgrowth (Fig. 2H).

Immunofluorescent (IF) staining of injured and non-injured IVD shows the presence of substance P and GAP43 in the injured IVD and absence of such in healthy IVDs (Fig. 2I). Similarly, IF of DRG showed relatively increased expression of GAP43 and CGRP as well as TRPV1 (Fig. 2J) in the injury matched DRGs compared to shams.

### TRIM29, ENO3, CAVIN1, RAB3B, and IHH identified as a potential biomarkers of degeneration

Assessment of transcriptomics (Fig. 3A-B, G) and proteomics (Fig. 3C-D, H) showed that injured and healthy IVD are significantly different. Interestingly, injured IVD samples tended to be more variable than healthy IVD samples in transcriptomics while proteomics showed the opposite trend (Fig. 3B, 3D). For proteomics, comparison of tissue and cell samples from injured and healthy IVD demonstrated that though disc healthy status was the primary determinant of sample clustering, sample type (cells vs. tissue) type also influenced sample similarity with healthy and injured tissue samples tended to be more similar than healthy and injured tissue cell samples (Fig. 3C-D). Furthermore, differentially expressed proteins (DEPs, Fig. 3E) and differentially expressed genes (DEGs, Fig. 3F) from RNA-seq and proteomics identified 4 overlapping markers that may be promising biomarker candidates. TRIM29, which is linked with DNA damage and immune response [98], was found to be downregulated in injured IVD in both transcriptomics and proteomics. ENO3 which is involved in metabolism [13], was found to be significantly downregulated injured IVD transcriptomics but upregulated in proteomics. Finally, CAVIN2 [14] and RAB3B [76; 97] which are associated with vesicular trafficking and signaling, were found in injured IVD to be downregulated at both the transcriptomic and proteomic level (Fig, 3G).

**Figure 3:**
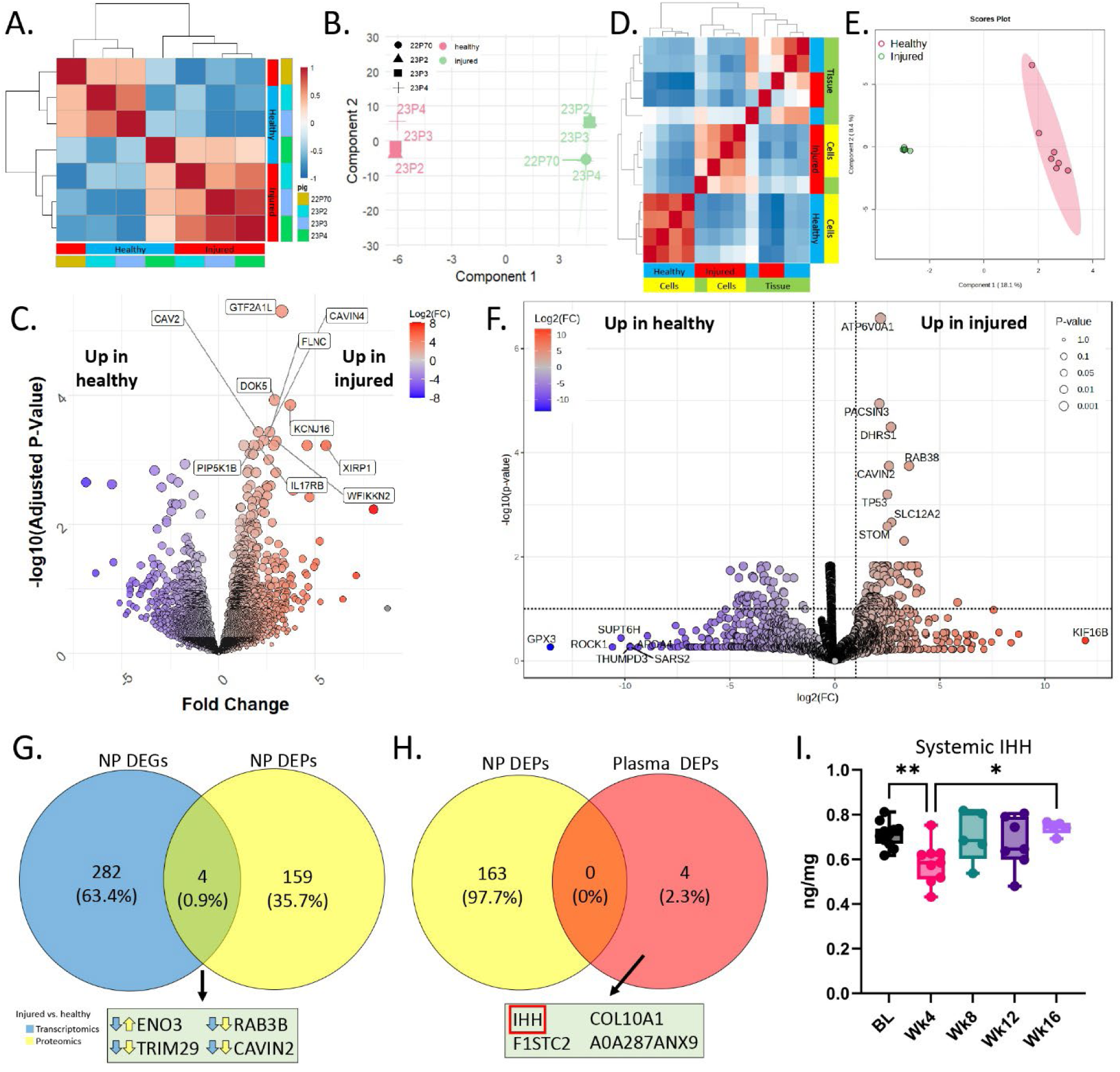
Proteomic and transcriptomic profiling reveals distinct, non-overlapping signatures and identifies circulating IHH as a potential biomarker. (A) correlation of all proteomic samples, (B) Sparse partial least squares analysis (SPLSA) of injured and non-injured IVD proteomics samples (C) correlation of all transcriptomic samples (D) (SPLSA) of injured and non-injured IVD transcriptomic samples, volcano plot of (E) proteomic DEPs and (F) transcriptomic DEGs; (G) overlapping DEP/DRGs, (H) Overlapping DEPs from IVD and plasma, (I) Systemic IHH concentrations.

In addition to IVD, proteomics was performed on plasma collected at baseline, week 12, and week 16 from the pigs throughout the experiment, which identified no overlapping DEP at either week 12 or 16 with IVD (Fig, 3H). Comparison on plasma from week 12 and 16 to baseline found moderate overlap in overall sample similarity as well as DEPs between samples (Supplemental fig 3).

Interestingly, Indian Hedgehog (IHH) was identified as a DEP in plasma at 12- and 16-weeks post-injury by proteomic analysis (Fig. 3H, supplemental fig. 3). Further quantification across all timepoint with ELISA revealed a significant decrease in systemic IHH concentration at 4-weeks post injury, followed by a gradual increase at later timepoints (Fig. 3I).

### Injury induces NC differentiation into NPC

Single cell transcriptomic analysis (scRNAseq) identified 21 clusters from healthy and injured porcine discs (Fig. 4A) collected at week 12 post-injury. Clusters were categorized as NC, NP, NC-NP, or EC) based on canonical markers expression as described in the methods (supplemental fig. 3). The prevalence of certain clusters the two types of discs (healthy vs injured) varied among the clusters, with some clusters such as NC2 primarily originating from healthy IVD while others like NP1 stemming primarily from injured IVD (Fig. 4B). Several clusters expressed markers from both NC and NP and thus were termed transitional NP (NC-NP).

**Figure 4:**
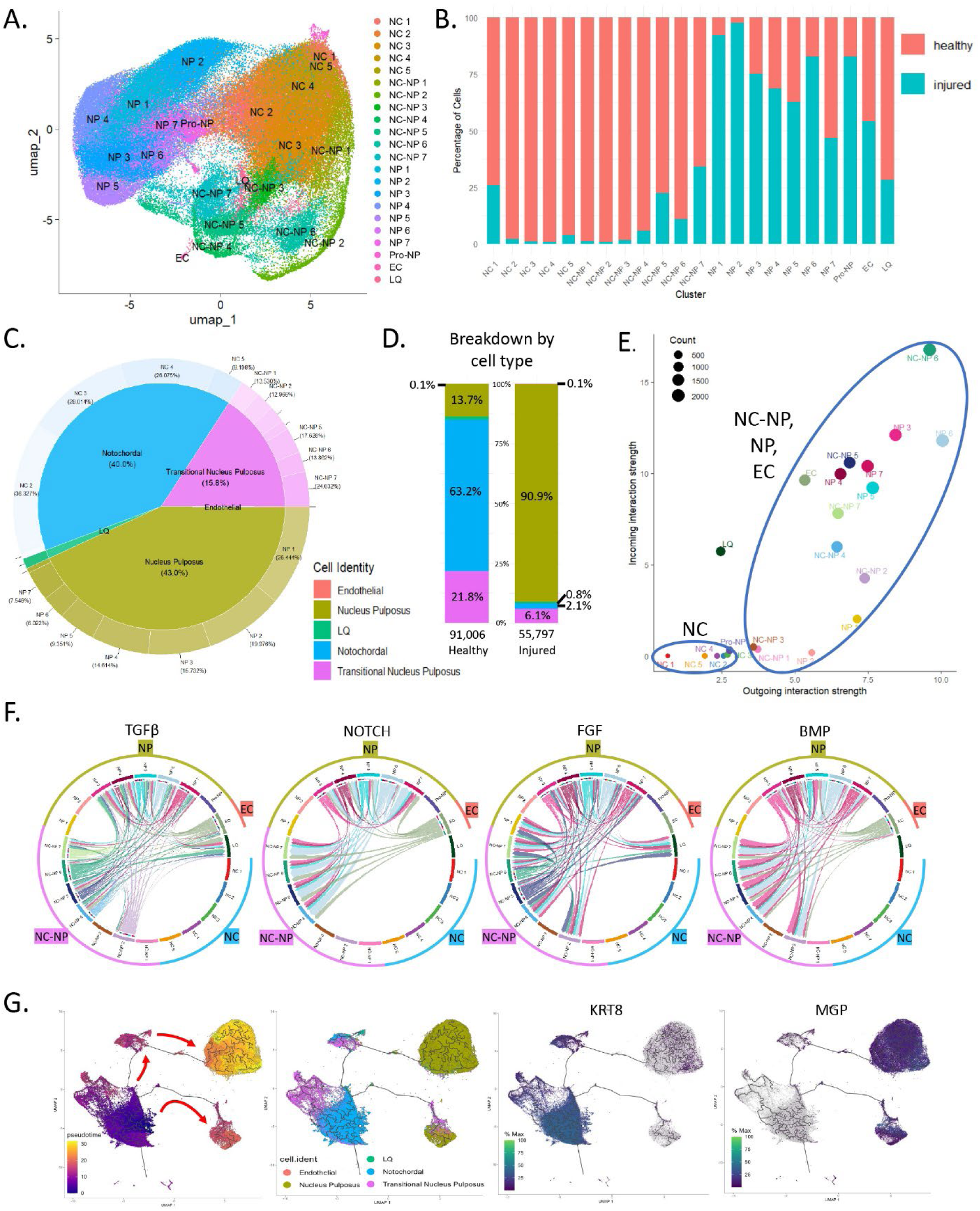
scRNA-seq demonstrates profound cell composition and signaling changes, suggesting differentiation of NCs in response to injury. (1) scRNA-seq UMAP colored and annotated by cluster, (B) proportional distribution of cell originating from healthy (red) and injured (blue) discs of each cluster, (C) summary of major cell identities and their major contribution across the dataset, (D) Comparison of cell type proportions between healthy (left) and injured IVD (right), (E) Relative outgoing and incoming intercellular communication strength of clusters, (F) Selected enriched intercellular communication pathways, (G) trajectory analysis from left to right colored by: pseudotime, cell type, KRT18 expression and MGP expression.

Closer examination of captured populations demonstrated that a majority of cells exhibited a NC (40.0%) or NP (43.0%) phenotype (Fig. 4C). However, when stratified by disc type, a majority of NCs were found to originate from the healthy IVD with 63.2% characterized as NCs compared to only 2.3% of cells from the injured IVD (Fig. 4D). In contrast, 90.9% of the cells captured from the injured IVD and only 13.6% of cells from the healthy IVD were identified as NP cells (Fig. 4D).

Analysis of intercellular communication via the CellChat package found that clusters characterized as NP or NC-NP have high intercellular communication compared to NCs (Fig. 4E). Pathways known to be associated with NC to NP differentiation such as TGFβ, Notch, BMP and FGF signaling were predicted to be enriched in NP clusters (Fig. 4F).

Trajectory analysis with Monocle 3 predicted that NCs tended to be located earlier in pseudotime compared to NP (Fig. 4G). Expression of characteristic markers of NC and IVD degeneration aligned with pseudotime results, with Keratin 18 (KRT18) found to be located earlier in pseudotime, while degenerative markers such as Collagen type 1A (COL1A1) were located later in pseudotime suggesting a transition from NCs to NPCs as a response to injury (Fig. 4I, Supplemental fig 4).

### Differentiated NPCs express increased pain inducing “help me” signaling

Interestingly, NPCs specifically were found to have comparatively increased expression of degenerative, inflammatory, and pain related markers compared to other cell types (Fig. 5A). CellChat analysis further identified multiple angiogenesis, inflammation, and neurogenesis/pain pathways that were differentially enriched in the NPC clusters such as VEGF [11; 75], NCAM [77], and prostaglandin [74; 90] signaling (Fig. 5B). IPA analysis of the NPC clusters identified that these cells were predicted to have high levels of senescence, cell stress, neural outgrowth, inflammatory signaling, leukocyte activation as well as various pathways associated with differentiation (Fig 5C). Comparison of the injured and healthy pig IVD with previously reported human LBP and asymptomatic IVDs [43] found that human LBP IVD were more similar to pig injured IVD than human asymptomatic IVD (Fig. 5D). Comparison of predicted enriched pathways from the NPCs of injured and healthy pig IVD as well as human LBP and asymptomatic IVD demonstrated that injured pig and human LBP samples were more enriched in cell stress and death and neural signaling pathways compared to healthy pig and human asymptomatic IVD (Fig. 5E). Conversely, healthy pig and human asymptomatic IVD were predicted to have greater enrichment in metabolic and stem cell/developmental pathways (Fig. 5E).

**Figure 5:**
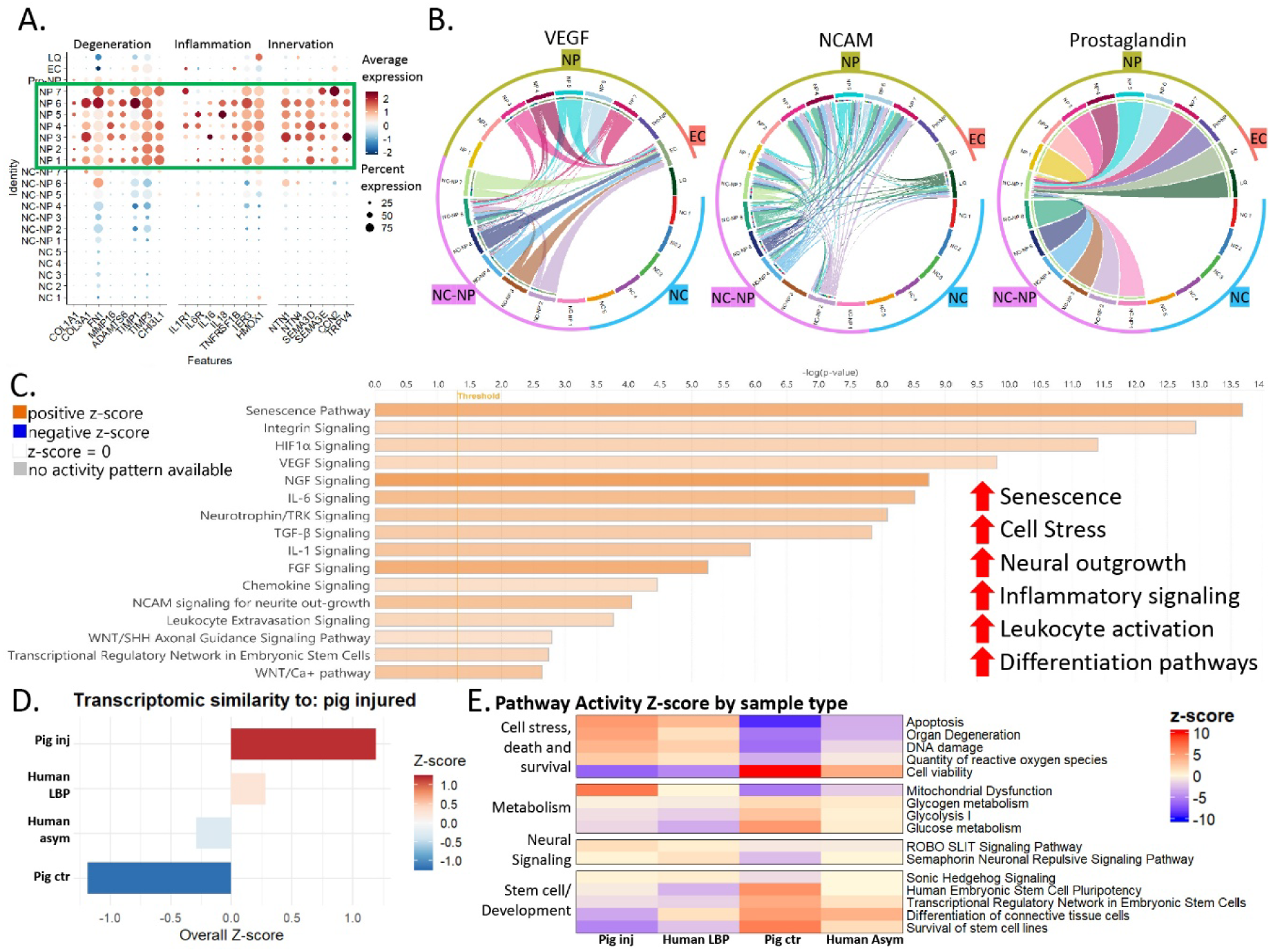
IVD injury results in the development of stressed NPCs which promote innervation and pain. (A) Dotplot of selected degeneration, inflammation and innervation markers; (B) Selected enriched vascularization, innervation, and inflammation pathways; (C) IPA pathway analysis of NPC clusters compared to other cell types; (D) Comparison of transcriptomic similarity by overall z-score to injured pig IVD. (E) IPA comparison of enriched pathways in NPCs from pig injured, pig healthy, human LBP and human asymptomatic IVD.

## Discussion

This study served as both a proof-of-concept and validation demonstrating that biobehavioral testing in pigs can be used to quantitatively assess the development of LPB. To date, there are no well-established large animal models of LBP with systematic behavioral assessments of pain. This work aims to fill this critical gap by establishing a translational model that combines a clinically relevant LBP model with quantifiable pain outcomes. In addition, this study characterizes the cellular and molecular changes that accompany disc injury, providing insight into the pathophysiological mechanisms contributing to pain (Fig. 6). These results were cross-referenced with previous findings in humans [43] therefore validating this model as clinically relevant for therapeutics development. Finally, we identified several candidate biomarkers for degeneration and LBP which, offers a valuable opportunity to investigate these biomarkers under controlled conditions.

**Figure 6:**
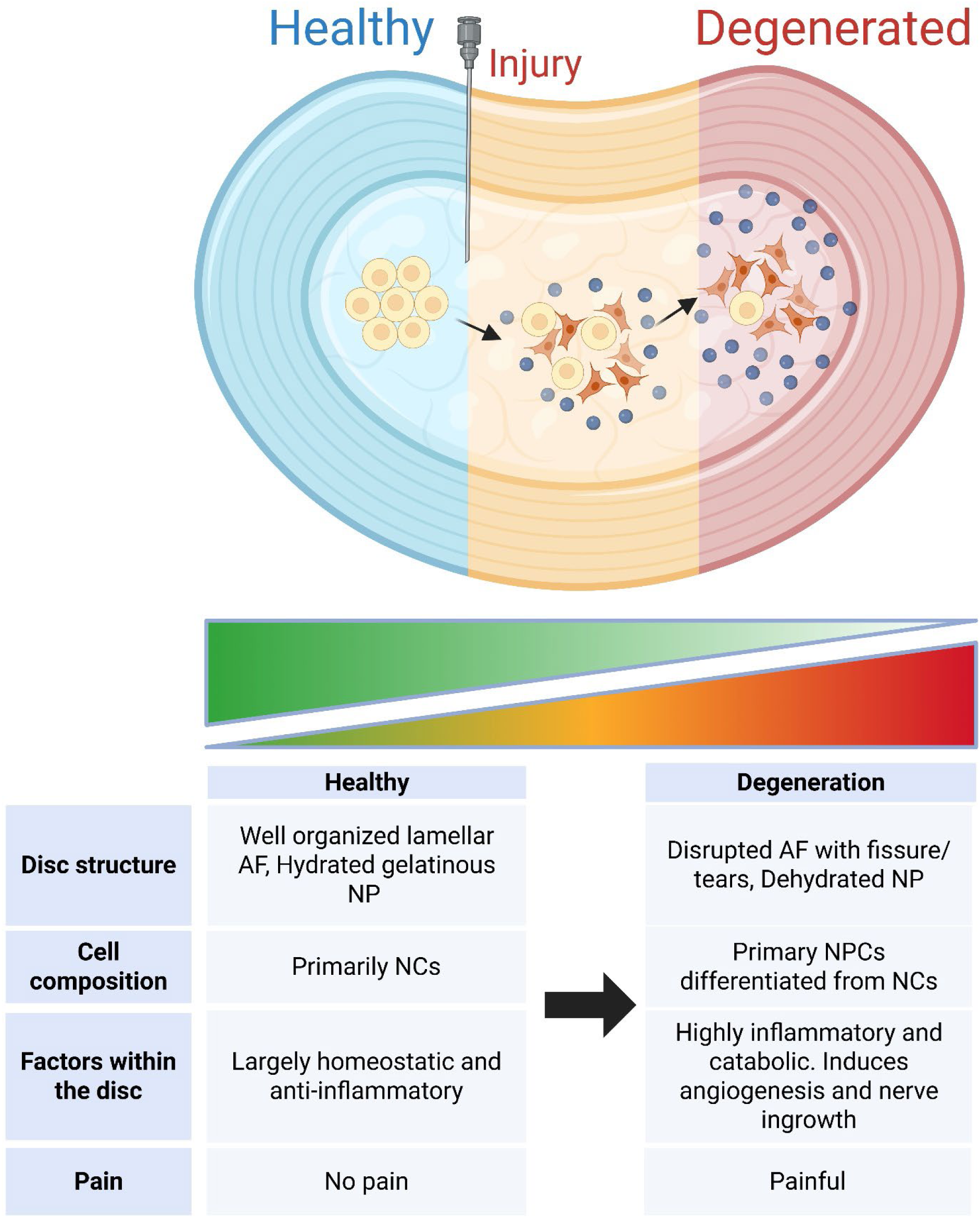
Proposed mechanism of disc degeneration and LBP development.

An important aim of this study was to develop reliable methods for assessing and quantifying pain-related behaviors in the porcine disc injury model. Biobehavioral testing was able to quantify and detect the pain responses observed in this model and was validated by qCEST (Fig. 2A) [56; 68], gene expression analysis (Fig. 2E) and transcriptomic analysis (Fig. 2F-G) of the DRGs. Furthermore, we were able to detect neuronal and nociceptive markers within the IVD itself via gene expression analysis (Fig. 1G) and IF (Fig. 2H), suggesting nerve ingrowth and increased nociceptive signaling within previously aneural disc tissue [10; 11; 26]. The ingrowth of nerves into the inner IVD has previously been observed in both human patients [10; 11; 28; 29] as well as other animal model [51; 82] suggesting that our disc injury model recapitulates a key pathological mechanisms of human IVD degeneration and LBP, thus making it a clinically relevant platform for translation of future LBP treatments.

Unlike humans who lose a majority of their NCs by age 10 [12], pigs retain their NCs well beyond skeletal maturity [95]. Thus, another goal of this study was to confirm that this model can faithfully recapitulate the cellular changes seen in human IVD degeneration. Our scRNA-seq revealed a near-complete loss of NCs and a corresponding shift towards NPCs following injury which closely mirrors the cellular landscape seen in degenerative human IVD (Fig. 4D). Notably, small population of NCs persisted in the injured IVD (Fig. 1B, 1D). This finding is consistent with our and others previously published work demonstrating that, contrary to the assumptions that NCs are completely lost post-maturity, some NCs are retained in the adult human IVD and can be detected even in individuals as old as 70 [44; 72; 93].

Beyond its translational value, our study also offers valuable insight into the cellular dynamics underlying IVD degeneration. Multiple lineage tracing studies have demonstrated progressive differentiation of NCs into NPCs with age and degeneration [3; 59; 83]. Our data aligns with these findings with trajectory analysis predicting that NC-rich healthy IVD are located earlier in pseudotime (Fig. 4G), indicating a less differentiated, progenitor-like cellular state compared to NPC-dominated injured IVD.

Furthermore, cell-cell communication analysis identified multiple pathways involved with NC-to-NPC differentiation in NC-NPC and NPC clusters, including Notch, TGFβ, BMP and FGF (Fig. 4F). Among these, the Notch and TGFβ pathway are particularly notable for their well-established and essential role in IVD development, growth, maintenance, and degeneration [45]. Notch regulates differentiation and tissue homeostasis by maintaining progenitor cells in an undifferentiated, proliferative state and directing their transition differentiated phenotypes [104]. It is essential for NC-to-NPC differentiation [73], elevated in degenerative musculoskeletal disorders including IVD degeneration [55; 104], and promotes NPC survival by inhibiting apoptosis [57]. Notch signaling is also influenced by proinflammatory cytokines such as IL1β and TNFα [91] [47], suggesting a potential link between inflammatory and cell fate within the IVD.

TGFβ is likewise critical for NC-to-NPC differentiation during development and postnatal IVD maintenance [18]. TGF-β promotes upregulation of NP markers expression, ECM synthesis, and phenotypic stability by enhancing proliferation and inhibiting apoptosis [53]. It also regulates cellular responses to mechanical and inflammatory stress, thereby contributing tissue homeostasis [9; 102]. However, dysregulated TGFβ signaling is also associated with fibrosis, aberrant matrix remodeling [106], and impaired regenerative potential [53]. Enrichment of these pathways support the hypothesis that injury induces NC-to-NPC differentiation and are consistent with developmental and degenerative transitions within the IVD.

Due to this close temporal and spatial relationship, there has been growing interest in studying NCs as both a key regulator of disc homeostasis and as a potential candidate for regenerative therapies. However, our understanding of NCs has been limited due to the limited availability of NCs in human tissue, thus making it difficult to study in clinically relevant contexts. In addition to advancing our understanding of IVD degeneration, elucidating the mechanisms governing NCs fate could inform the development of NC based therapies for IVD degeneration and discogenic LBP.

In addition to exploring the NC niche, we demonstrated the presence of highly transcriptionally active NPCs which we had previously observed and described in human LBP samples [43]. Our scRNA-seq demonstrated similar upregulation degeneration-, inflammation-, and innervation-associated markers in NC-NPC and NP clusters (Fig. 5A). Further analysis of these clusters via CellChat and IPA demonstrated enrichment of signaling pathways related to innervation, vascularization, and inflammatory pathways (Fig. 5B-C). To evaluate translational relevance of our model, we compared the transcriptomic profiles of our pig IVD with our previously published human asymptomatic and LBP IVD. This analysis revealed injured pig IVD were more transcriptionally similar to human LBP IVD than human asymptomatic IVD (Fig. 5D). Moreover, pathways associated with cellular stress, altered metabolism, neural signaling, and stem cell development were consistently stratified between the injured/LBP and healthy/asymptomatic IVDs. This suggests that injury-induced changes in the porcine IVD recapitulate not only the cellular alterations but also the molecular pain-associated phenotype observed in human discogenic low back pain. These findings reinforce the suitability of this model for studying both IVD degeneration and discogenic LBP.

We identified several differentially expressed biomarkers of interest which are promising avenues of future study. Proteomic and transcriptomic comparison of upregulated markers identified four markers that we significant at both levels of expression: ENO3, RAB3B, CAVIN2, and TRIM29. In particular, RAB3B and CAVIN2 are markers of caveolar and vesicular trafficking and have been previously linked to cellular signaling regulation, membrane dynamics, and mechano-transduction [65; 80]. Although RAB3B has previously been reported to be downregulated in degenerating IVD [21] and CAVIN2 has been studied as a target to enhance extracellular vesicle uptake in NPCs [54], their potential roles in LBP pathogenesis have not, to our knowledge, been explored. Interestingly, examination of cellular communication and signaling mechanisms within the IVD using scRNA-seq revealed that NC clusters, which are prominent in healthy IVD, highly expressed genes associated with caveolae-mediated signaling (Supplemental Fig. 5). In contrast, NP clusters, which are predominately in the injured IVD, showed increased expression of genes involved in constitutive signaling and secretion, with marked downregulation of caveolae signaling genes (Supplemental Fig. 5). These findings demonstrate, not only that RAB3B and CAVIN2 are promising biomarkers for IVD degeneration and potentially LBP, but also that cellular communication and signaling mechanisms change with degeneration. Notably, there is a lack of comprehensive studies examining how cell–cell communication networks shift in the context of IVD degeneration and LBP making it a promising area for future study.

In addition, to IVD local markers, we also identified IHH as a plasma DEP at 12- and 16-weeks post-injury. IHH, which is well known for its critical role in IVD development and homeostasis, plays an essential role in NC maintenance and the structural organization of the developing IVD [19]. It is typically highly expressed in healthy, non-degenerated discs; however, several studies have shown that loss of IHH expression is associated with pathological transitions, including the loss of NCs and the emergence of chondrocyte-like NPCs [4; 103]. Notably, a study by Bach et al. in a canine model reported the biphasic pattern of IHH expression observed in our proteomics and ELISA results, namely the high IHH in healthy IVD, reduction with the loss of NCs, and subsequent increase as IVD degeneration progressed to later stages of degeneration [4]. These results suggest that IHH may be a potential biomarker for the early-to-intermediate stage of IVD degeneration.

There are several limitations to this study. First, we used intra-pig non-injured controls for several of our experiments, including bulk and scRNA-seq, histology, and proteomics, which has the advantage of using the same animal with the same genetic background, but may also be a limitation due to some systemic effect of the injury that could not be completely excluded. Furthermore, we did not analyze any earlier timepoints for our transcriptomic analyses, which limits our understanding of the important initial molecular and cellular changes during early disc degeneration. Finally, due to the relative scarcity of pig-specific bioinformatic resources in biomedical research, many of the databases, such as CellChat and IPA, were analyzed using existing human settings, which may affect the accuracy of these analyses.

To assess structural contributors to pain, adjacent vertebrae were evaluated for Modic changes, which are present in 18-58% of LBP cases [1]. Type I and, to a lesser extent, Type II Modic changes are most commonly associated with pain, due inflammation and sensitization [1]. In our model, though Type I changes appeared at earlier timepoints, acute pain was not detected until week 10, when most vertebrae exhibited Type II or III changes. This suggesting that factors beyond Modic changes may contribute to LBP. However, distinguishing discogenic from vertebrogenic pain remains challenging due to the anatomical and functional integration of the IVD and adjacent vertebra.

This study employed two behavioral pain assessments, the Glasgow Composite Pain Scale and the wind-up ratio test, which specifically evaluate acute nociceptive responses and temporal summation, respectively. While these assays provided valuable information on pain sensitivity and central sensitization, they primarily capture evoked pain responses and do not fully represent the complexity of pain IVD injury may induce. A more comprehensive evaluation of discogenic pain would benefit from the inclusion of additional assessments targeting spontaneous behaviors, affective components of pain, and longer-term functional outcomes.

## Conclusion

In conclusion, our study demonstrates the potential use of a combined novel MRI and biobehavioral testing to quantify pain in a porcine IVD injury model of discogenic LBP and suggests using TRIM29, ENO3, RAB3B, CAVIN2, and IHH as potential biomarkers for LBP. Furthermore, this study demonstrates the appropriateness of this model for translation of LBP therapies towards clinical adoption as it closely mirrors the degenerative phenotype and symptoms seen in human discogenic pain.

## Data Availability

The sequencing data generated and analyzed during this study have been deposited in the Gene Expression Omnibus (GEO) under accession number GSE308921. All code used for data processing and analysis is available at: https://github.com/gkaneda/Pig-IVD-Scripts.

## Acknowledgements

We would like to thank Ed Paredes and Adrian Glenn for assisting us with fluoroscopic imaging during injury induction. We would also like to thank Dr. Eugenio Cingolani for generously providing us with sham IVD and DRG tissues from the pigs used in his studies. This work was supported in part by the Cedars-Sinai Applied Genomics, Computational and Translational Core, the Cedars-Sinai Research Imaging Core, the Cedars-Sinai Proteomics and Metabolomics Core, and the Cedars-Sinai Biobank and Research Pathology Resource.

## Graphical Abstract

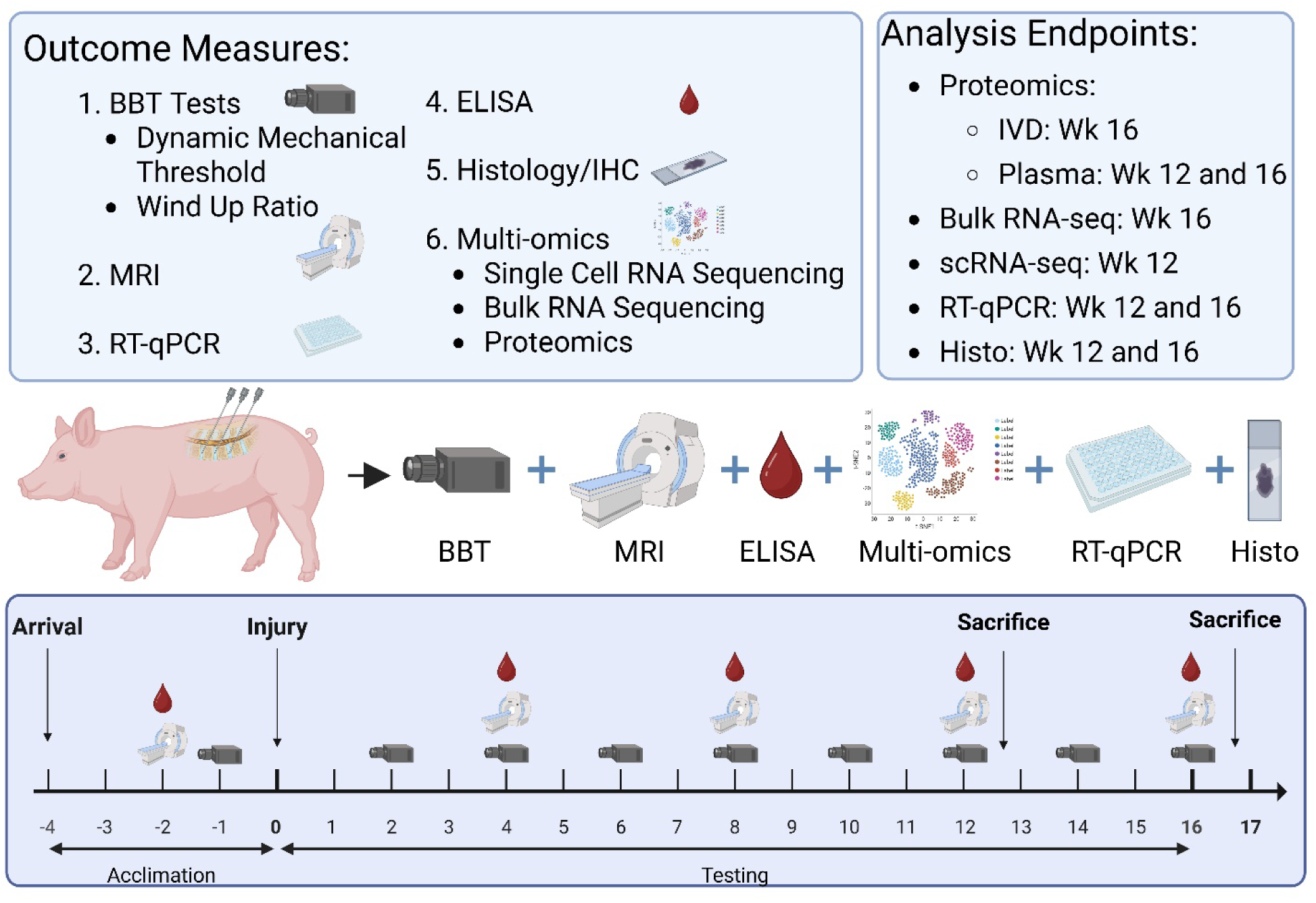

