## Supplemental Figures and Tables for "A Porcine Model of Intervertebral Disc Injury Recapitulates Human Discogenic Pain via Notochordal Cell Loss and Pain-inducing Nucleus Pulposus Cell Emergence"

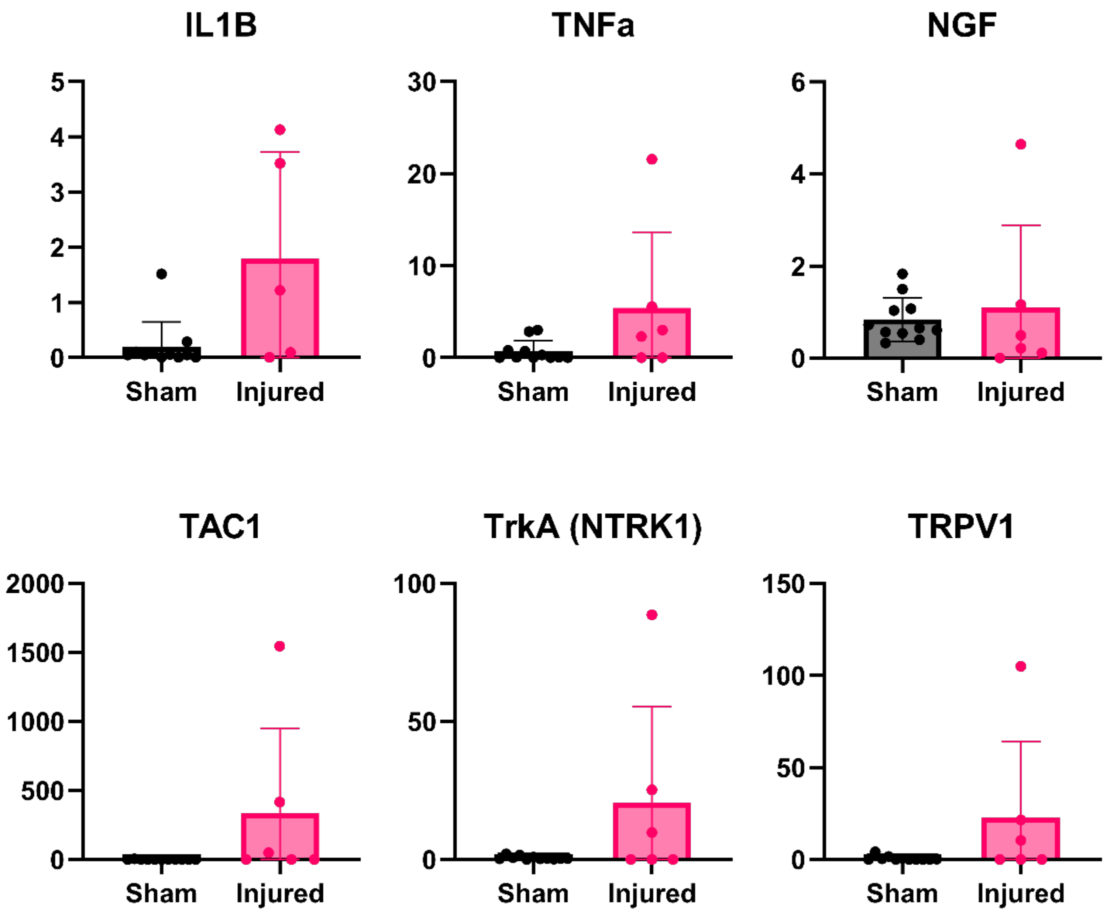


Supplemental Figure 1: Gene expression for inflammatory (IL-1β, TNFα), NGF related (NGF, TrkA) and nociceptive/peptide channel (TAC1, TRPV1) genes in injured and sham IVD.

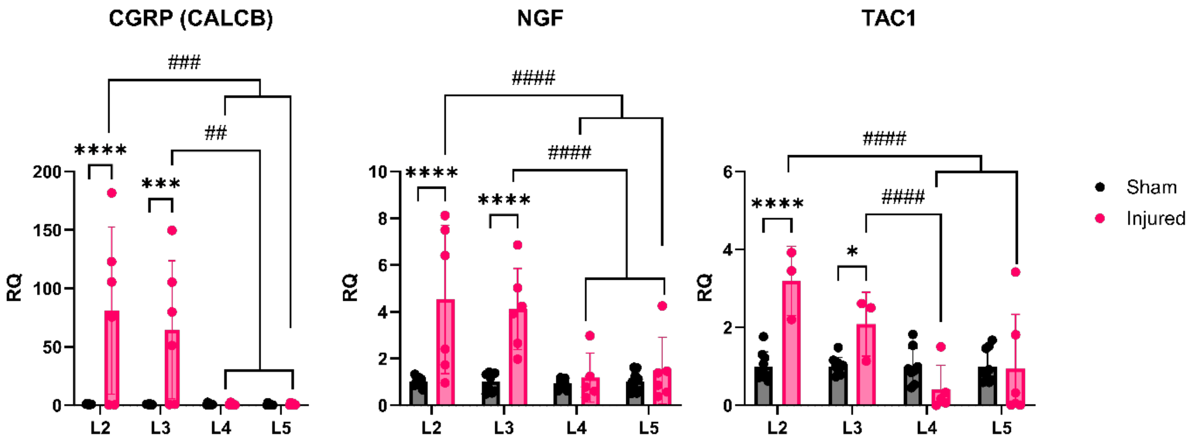


Supplemental Fig. 2: Comparison of genes across level and injury status. *** p<0.001, ****p<0.0001 between groups at the same level. ## p<0.01, ### p<0.001, #### p<0.0001 between levels in the injured group.

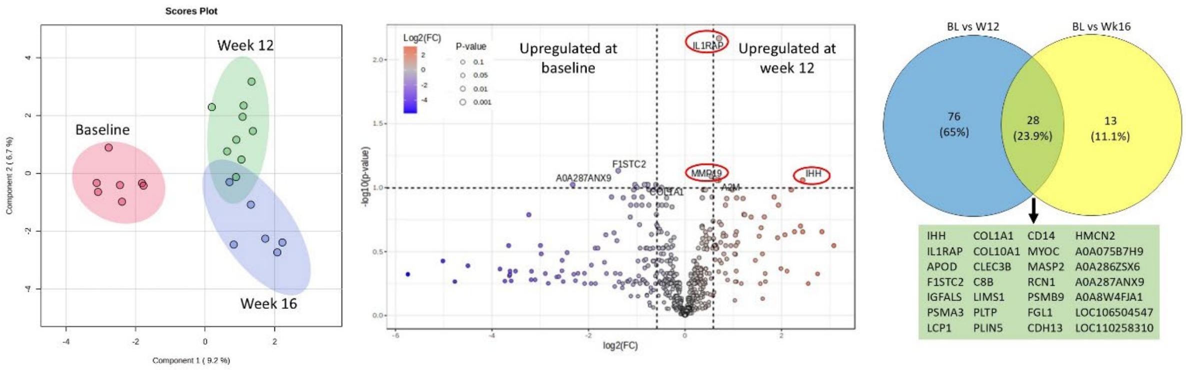


Supplemental fig. 3: Comparison of plasma proteomics at BL, week 12 and week 16. DEP cut-off was set to raw p-value < 0.05. Left: sPLS of baseline, week 12 and week 16 samples. Middle: Volcano plot of BL vs week 12 DEPs. Right: Overlapping DEPs identified at week 12 and week 16.

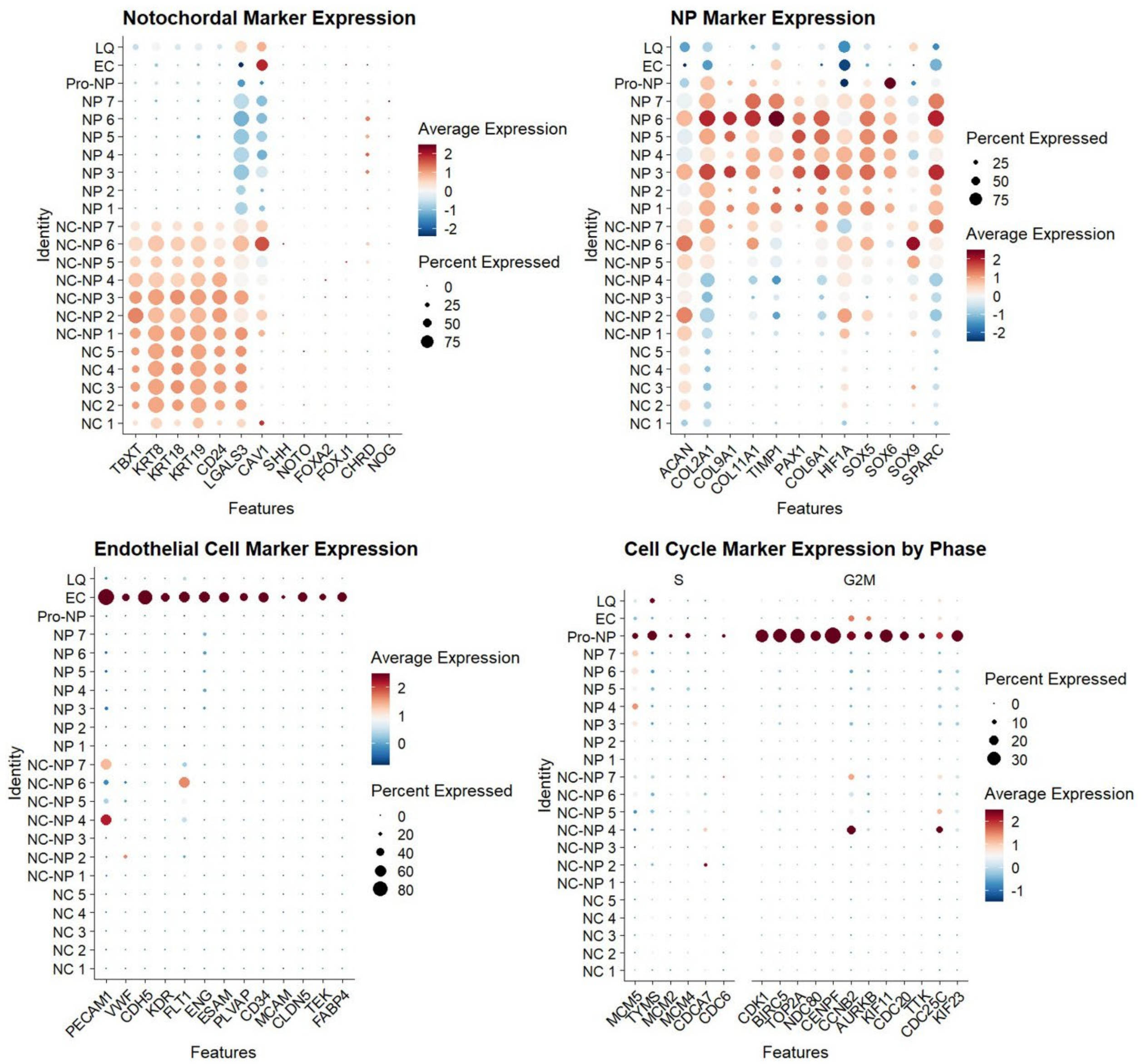


Supplemental Figure 4: Dotplot of canonical genes used to identify cell identities. Top left: NC; Top right: NP; Bottom left: EC; and Bottom right: proliferating NP.

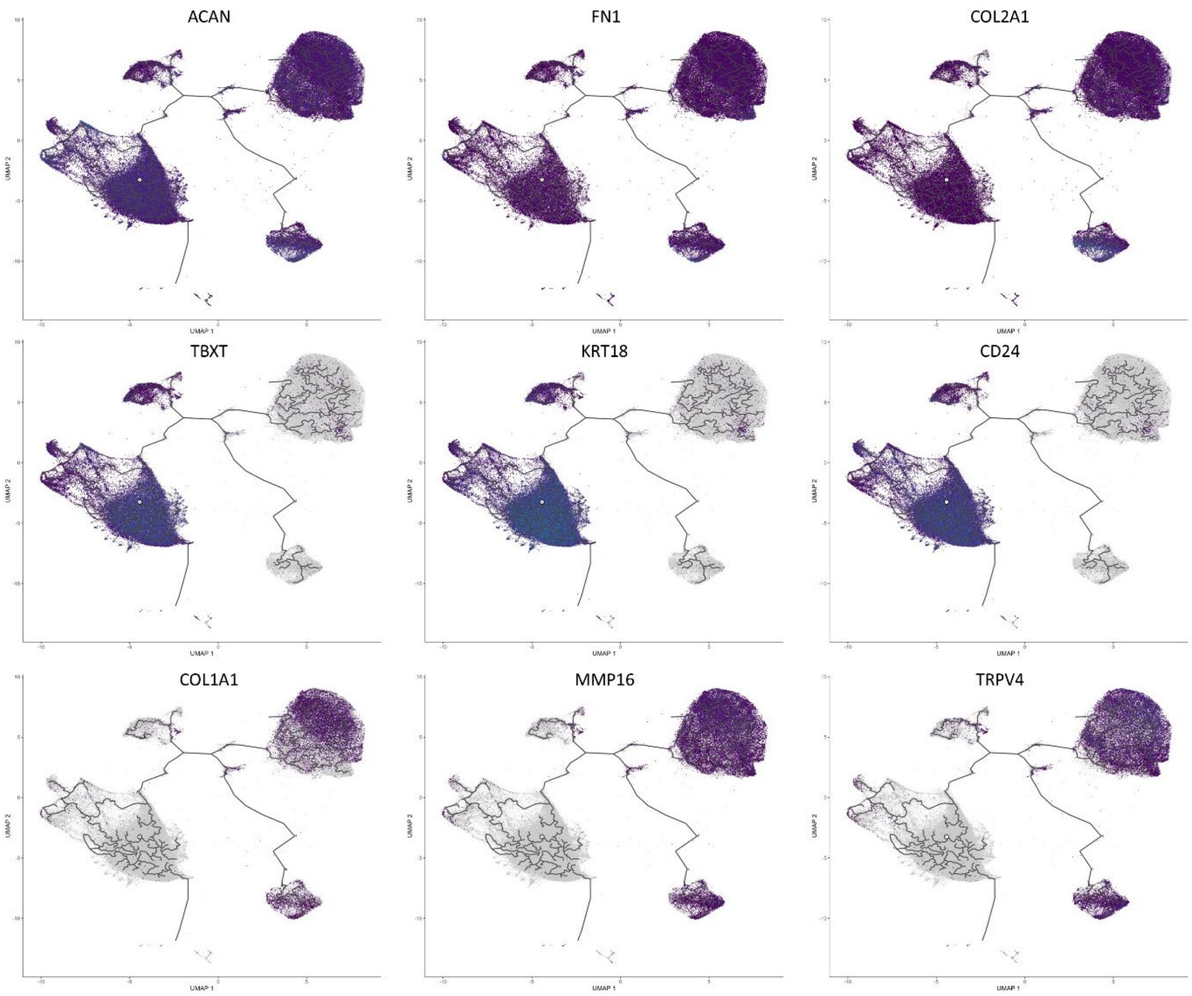


Supplemental Figure 5: Expression of Canonical IVD (top row), notochordal (middle row), and degenerative markers along pseudotime (bottom row).

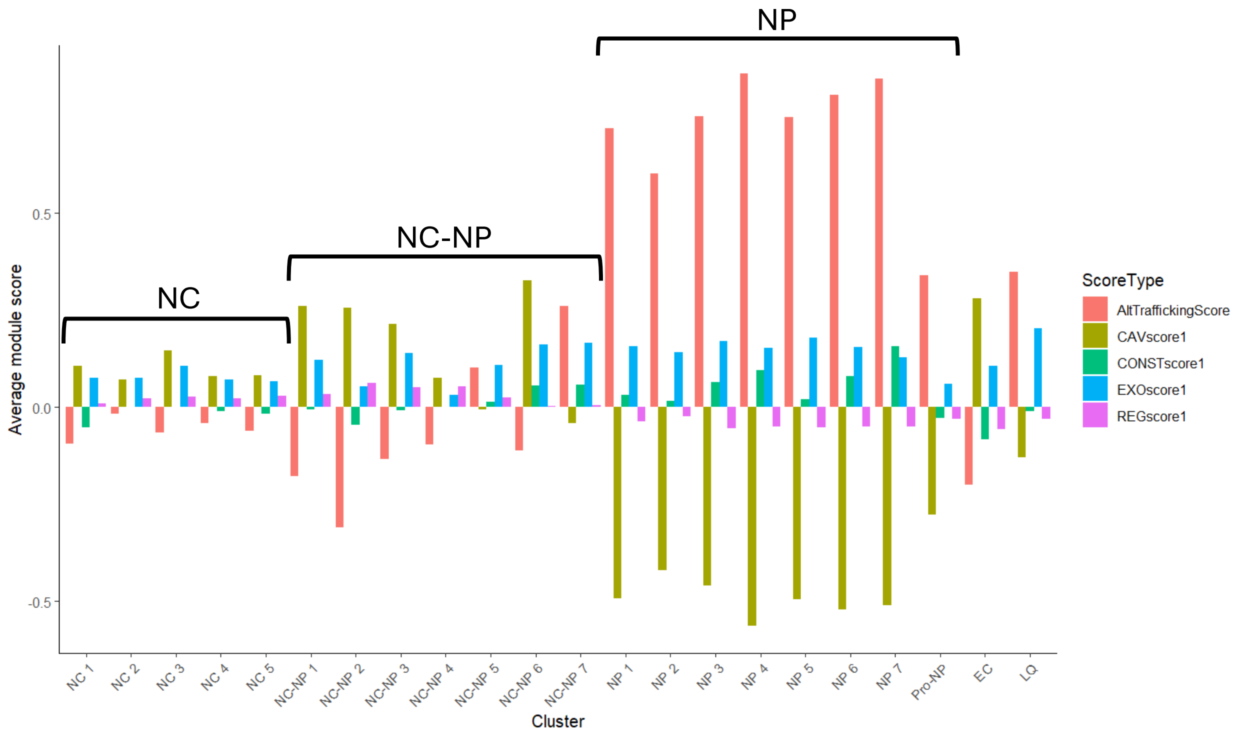


Supplemental fig 6: Average module scores for caveolae/caveolin signaling (CAVscore1), constitutive secretion (CONSTscore1), Exosome/microvesicle (EXOscore1), regulated/synaptic-like exocytosis (REGscore1), and Composite score (AltTraffickingScore) comparing exosome/constitutive routes (positive score) to caveolae/regulated routes (negative score). AltTraffickingScore = (EXOscore + CONSTscore) − (CAVscore + REGscore).

Supplemental Table 1: Modified Glasgow Pain Scale Scoring Criteria

| Comfort | | Score |
|---|---|---|
| | Awake, interested in surroundings, pig recumbent or standing, eating pellets when offered, vocalizations (grunts, barks). | 0 |
| | Awake, standing or recumbent, not interested in surroundings, eating pellets when offered, vocalizations (grunts, barks, squeals). | 1 |
| | Lethargic, depressed appearance, reduced appetite for pellets, eating treats as last resort for behavior tasks, vocalizations (grunts, barks, squeals). | 2 |
| | Prone/recumbent, lethargic, no appetite for pellets, reduced appetite for treats, unwilling to perform behavior tasks, vocalizations (strained, screams, absence of). | 3 |
| | Lateral, lethargic, depressed mentation, no appetite, reluctant to rouse or move, fixed look & staring /or eyes half closed, vocalizations (strained scream, absence of). | 4 |
| **Social Behavior** | | **Score** |
| | Normal, shows robust interest in investigator, interacts with adjacent housed pigs or pen-mates. | 0 |
| | Mild changes, moderate interest in investigator, slower to interact, little to no interest in adjacent housed pigs or pen-mates. | 1 |
| | Moderate changes, little to no interest in investigator, decreased responses on approaching or interactions, no interest in adjacent housed pigs. | 2 |
| | Severe changes, no interest or response to investigator or other pigs. | 3 |
| **Surgical or Target Site** | | **Score** |
| | Ignores target site. | 0 |
| | Consistent rubbing of target site, by self or on housing. | 1 |
| **Movement** | | **Score** |
| | Normal ambulation, full weight-bearing, no lameness. | 0 |
| | Slight lameness (limping, toe touching) due to target site sensitivity. | 1 |
| | Lameness, limping / toe touching on some, but not all steps. | 2 |
| | Lameness, limping/toe touching on all steps when walking voluntarily. | 3 |
| | Refusing/unable to stand, without or without assistance, remaining recumbent or lateral. | 4 |
| **Response to Site Palpation** | | **Score** |
| | No observable change in behavior. | 0 |
| | Looks around. | 1 |
| | Flinches, shies away, vocalizes (barks, squeals). | 2 |
| | Vocalizes (scream), tries to escape. | 3 |

Supplemental Table 2: Taqman primers information

| Gene | Assay ID | PubMed RefSeq |
|---|---|---|
| MMP13 | Ss03373279_m1 | XM_003129808.5 |
| IL-1β | Ss03393804_m1 | NM_214259.2 |
| TNFa | Ss03391318_g1 | NM_214022.1 |
| PGP9.5 (UCHL1) | Ss03381306_u1 | NM_213763.2 |
| CALCRL | Ss03394018_m1 | NM_214095.1 |
| CGRP (CALCB) | Ss03386461_u1 | NM_001104955.1 |
| RAMP1 | Ss06942043_m1 | NM_214199.2 |
| TRPV1 | Ss03377173_u1 | XM_013981216.2 |
| TAC1 | Ss04330568_gH | XM_003130163.6 |
| TrkA (NTRK1) | Ss06935350_g1 | XM_001929525.5 |
| NGF | Ss04330572_gH | XM_021089996.1 |
| GFAP | Ss03373547_m1 | NM_001244397.1 |
| ATF3 | Ss06896040_m1 | XM_003482749.4 |

Supplemental Table 3: Antibodies

| Target | Host | Manufacturer | Cat # |
|---|---|---|---|
| COL2 | Mouse | Novus Biologicals | NBP1-05169 |
| GAP43 | Rabbit | Novus Biologicals | NB300-143 |
| SubP (IVD) | Rabbit | Sigma Aldrich | S1542 |
| SubP (DRGs) | Rat | Novus Biologicals | NB100-65219 |
| CGRP | Goat | LsBio | LS-C41796-0.1 |
| NGF | Mouse | MyBioSource | MBS2138403 |
| PGP9.5 | Mouse | Invitrogen | 480012 |
| TRPV1 | Rabbit | Novus Biologicals | NB100-98897 |
